# Impaired expression of chloroplast HSP90C chaperone activates plant defense responses leading to a disease symptom-like phenotype

**DOI:** 10.1101/2020.04.07.029116

**Authors:** Islam Shaikhul, Bhor Sachin Ashok, Tanaka Keisuke, Sakamoto Hikaru, Yaeno Takashi, Kaya Hidetaka, Kobayashi Kappei

## Abstract

RNA-seq analysis of a transgenic tobacco plant, i-hpHSP90C, in which chloroplast *HSP90C* genes can be silenced in an artificially inducible manner resulting in the development of chlorosis, revealed the up- and down-regulation of 2746 and 3490 genes, respectively. Gene Ontology analysis of these differentially expressed genes indicated the upregulation of ROS-responsive genes, the activation of the innate immunity and cell death pathways, and the downregulation of genes involved in photosynthesis, plastid organization, and cell cycle. Cell death was confirmed by trypan blue staining and electrolyte leakage assay and the H_2_O_2_ production by diaminobenzidine staining. The upregulation of ER stress-responsive genes suggested the interplay between ER protein quality control and chloroplast or immune response. The results collectively suggest that the reduced levels of HSP90C chaperone leads the plant to develop chlorosis primarily through the global downregulation of chloroplast and photosynthesis-related genes and additionally through the light-dependent production of ROS, followed by the activation of immune responses including the cell death.

**Highlight:** Induced silencing of *HSP90C* gene caused the upregulation of stress-responsive genes and the activation of innate immune response, which resulted in the chlorosis development accompanying cell death.

## Introduction

Plant virus diseases develop a range of symptoms resulting from the morphological and physiological disturbances of the host cells. Leaf chlorosis, the most frequently observed symptom, reduces plant productivity and thus, leads to a significant loss in crop yield. Virus-induced chlorosis often accompanies the structural change and dysfunction of chloroplasts, including the reduction in chlorophyll content and the expression of photosynthetic genes (Manfre *et al*., 2011; Mochizuki and Ohki, 2011; Mochizuki *et al*., 2014; Qiu *et al*., 2018). Therefore, understanding the mechanisms of chloroplast dysfunction would lead us to the establishment of counter-measures against crop loss. Although the molecular events during chlorosis have been extensively documented, the precise mechanism for the reduced chloroplast activities had remained to be elucidated until recent pioneering studies, which have shown the involvement of RNA silencing of chloroplast protein genes in the development of chlorosis by sub-viral RNAs. Two groups have independently shown that the bright yellow symptoms in tobacco plants infected with cucumber mosaic virus (CMV) harboring Y-satellite RNA (Y-sat) is attributed to the RNA silencing of magnesium protoporphyrin chelatase subunit I (CHLI) involved in chlorophyll biosynthesis mediated by the Y-sat-derived small interfering RNA (siRNA) (Shimura *et al*., 2011; Smith *et al*., 2011). Another study has shown that siRNA derived from *Peach latent mosaic viroid* (PLMVd) directs the RNA silencing of chloroplast heat-shock protein 90 (HSP90C) and consequently causes severe chlorosis or albinism (Navarro *et al*., 2012). To analyze the molecular mechanisms of chlorosis induced by the RNA silencing of chloroplast proteins, we previously established experimental systems using chemically inducible promoter to drive RNAs to induce RNA silencing of CHLI and HSP90C in transgenic tobacco plants (Bhor *et al*., 2017*a*,*b*).

Heat shock proteins or HSPs are a group of multiple protein families, which are induced upon different cellular stresses and help cells survive under stressed condition (Parsell and Lindquist, 1994). HSPs are evolutionarily conserved throughout the prokaryotes and eukaryotes and have been classified into different families, including HSP100, HSP90, HSP70, HSP60, HSP40, and HSP20 (Park and Seo, 2015; Sable *et al*., 2018). All of these families help maintain cellular homeostasis and play roles in different developmental processes through their molecular chaperone function (Park and Seo, 2015; Sable *et al*., 2018). Out of these families, HSP90 family proteins have been shown to play pleiotropic roles in diverse biological phenomena. HSP90 null mutants are known to be lethal, and even the heterozygotes for the mutation showed developmental abnormalities in *Drosophila* (Rutherford and Lindquist, 1998). The pharmacological blockade of HSP90 function resulted in phenotypic variations specific to genetic backgrounds (Queitsch *et al*., 2002). The studies support that HSP90 family proteins take wide-ranging client proteins for their chaperoning function in different signal transduction pathways and transcriptional regulatory networks (Echeverría *et al*., 2011; Wayne *et al*., 2011).

In plants, the HSP90 family comprises 7, 9, and 10 members (Krishna and Gloor, 2001; Mandal *et al*., 2013) in *Arabidopsis thaliana, Oryza sativa*, and *Populus trichocarpa*, respectively (Zhang *et al*., 2015). Out of seven *Arabidopsis* HSP90 family proteins, four members exhibit nucleo-cytoplasmic localization, while the other three localize to each of chloroplast, mitochondrion, and endoplasmic reticulum (Krishna and Gloor, 2001). The HSP90C or AtHSP90.5, which localizes to the chloroplast, was first identified as the causal gene for a chlorate-resistant mutation (Cao *et al*., 2003). Like cognate proteins in the mitochondrion (Altieri *et al*., 2012; Altieri, 2013) or endoplasmic reticulum (Ishiguro *et al*., 2002; Marzec *et al*., 2012), HSP90C has been proposed to have a role in protein folding in the chloroplast (Schroda and Mühlhaus, 2009) and shown to be essential for the transport of different proteins into chloroplasts (Inoue *et al*., 2013). Although the chlorate-resistant mutant, *cr88*, which has a single missense mutation in HSP90C coding sequence, was viable albeit with an impaired photomorphogenesis (Cao *et al*., 2003), null mutants with T-DNA insertions in HSP90C gene were lethal (Inoue *et al*., 2013; Feng *et al*., 2014), indicating its essential function. The silencing of HSP90C in Arabidopsis plants resulted in variegated or albino phenotype (Oh *et al*., 2014). Therefore, it is not surprising that the silencing of HSP90C by viroid-derived siRNA resulted in severe chlorosis or albinism (Navarro *et al*., 2012).

We previously showed that the induced silencing of HSP90C in transgenic tobacco resulted not only in visible chlorosis with significantly decreased chlorophyll contents and the reduced expression of chloroplast protein genes but also in the induction of some pathogenesis-related genes (Bhor *et al*., 2017*b*). The results suggest that the chlorosis in this model system is attributed not only to the impaired chloroplast biogenesis but also to active plant responses to the impaired chloroplast function. In this report, we employed an RNA-seq to examine the alteration in gene expression early after the induction of HSP90C silencing. By comparing the gene expression patterns between HSP90C silenced and non-silenced plants, we found upregulation of genes related to the response to reactive oxygen species, cell death, plant hormone signaling pathways, defense response, and innate immune response. The detection of sporadic cell death supported the biological significance of the transcriptomic changes. The results suggest that chlorosis development with impaired HSP90C expression involves the activation of cell-death mediated plant defense response in addition to a simple reduction of chloroplast function.

## Materials and methods

### Plant materials

Transgenic tobacco lines, i-hpHSP90C 6-1 (H-6) and i-hpHSP90C 4-5 (H-4), which express hp-RNA (hairpin RNA) corresponding to the HSP90C-specific regions under the control of a dexamethasone-inducible promoter, were described previously (Bhor *et al*., 2017*b*). These inducible HSP90C silencing tobacco lines and non-transformant tobacco (*Nicotiana tabacum* cv. Petit Havana SR1) were used in this study. Plants were cultured in a plug tray containing commercial soil mix (Supermix A, Sakata Seeds, Yokohama, Japan) for one week at 25°C under 16/8 h light/dark cycle condition with an irradiation dose of about 60 μMm^−2^s^−1^. The seedlings were transferred and grown for additional 2 weeks in a plastic pot (6.0 cm in diameter) containing the mixture (1:1) of vermiculite and Supermix A, with watering every other day with 1000-times diluted Hyponex 6-10-5 (Hyponex Japan, Osaka, Japan) solution twice a week. To induce the transgene expression, three-week-old control and transgenic tobacco plants were sprayed with freshly diluted 50 μM dexamethasone (Dex) solution containing 0.01% (v/v) Tween-20 using a spray bottle. Control plants were mock-treated by spraying with 0.5% ethanol solution containing 0.01% (v/v) Tween-20.

### Extraction and sequencing of RNA

The RNA was extracted and treated with RNase-free DNase as described previously (Waliullah *et al*., 2014). For RNA-seq analysis, RNA was extracted from six individual plants each Dex-treated and untreated, control and H-4 (i-hpHSP90C 4-5) transgenic plants at 24 h post-Dex treatment. Three samples each from plant/treatment groups were selected by the integrity of RNA assessed using 2100 Bioanalyzer with RNA-nano chip (Agilent). Libraries were constructed using the TruSeq RNA Sample Preparation v2 kit (Illumina) according to the manufacturer’s protocol. 100-bp single-end sequencing was carried out using Hiseq2500 at Nodai Genome Research Center.

### RNA-seq data analysis

Raw reads were obtained using bcl2fastq2 (Illumina) with the adaptor sequences removed. The read data were further trimmed using fastq_quality_trimmer with the quality cutoff at 28 and length cutoff at 80. The clean read data were uploaded to the local Galaxy platform (Boekel *et al*., 2015), and analyzed therein using Salmon (Patro *et al*., 2017) with Ntab-TN90-AYMY-SS_NGS.mrna.annot.fna reference transcriptome (Sierro *et al*., 2014); downloaded from Sol Genomics Network; https://solgenomics.net/organism/Nicotiana_tabacum/genome) for transcript abundance estimation and expression value computation. We prepared in the command line from the fasta file above to use a Transcript ID-AGI code table (Supplementary table S1) to have the expression values with AGI codes because the original gene IDs in the accompanying gff3 file were not compatible with our GO enrichment analysis. Thus, Salmon gave the Transcripts Per Million (TPM) value to each of the transcripts with AGI code annotation. Of 189,413 mRNA transcripts in tobacco reference transcriptome, 121,268 transcripts (64.02%) were annotated with AGI codes, and 68,145 transcripts (35.98%) were omitted from the analysis (Supplementary table S1).

The detection of differentially expressed genes (DEGs) was performed using DESeq2 (Anders and Huber, 2010) in the Galaxy platform. DESeq2 was run in different groupings: Dex-treated H-4 vs control H-4; Dex-treated H-4 vs Dex-treated SR1; and Dex-treated H-4 vs Control SR1. Then, Venn diagrams showing the DEGs in various combinations were prepared using the web program Venny 2.1 (http://bioinfogp.cnb.csic.es/tools/venny/index.html). The differentially expressed genes were considered significant under the following criteria: corrected p-value (P(adj)) less than 0.05 and the Log_2_(FC) values above 1 or below −1. For functional annotation of DEGs with AGI codes, we performed the GO enrichment analysis using the PANTHER (Protein ANalysis THrough Evolutionary Relationships) Classification System (http://www.pantherdb.org/) (Mi *et al*., 2019), which implements one-sided Fisher’s exact test with the multiple-testing correction method being set to FDR. GOs with FDR below 0.05 were considered significant. Heatmap for selected up-regulated genes was drawn using ggplot2 function in the heatmap2 program in the Galaxy platform. The relative normalized count data calculated in the Excel program (Microsoft) were used instead of the raw normalized count data from DEseq2 because the latters were not accepted by the program. The expression levels of particular genes in the RNA-seq analysis were compared using the relative normalized count data with statistical analysis using DESeq2.

### Quantitative RT-PCR

Total RNA was extracted using the ISOSPIN Plant RNA Kit (Nippon Gene, Japan) for qRT-PCR with more plant samples. The cDNA was synthesized using the M-MLV RTase (New England Biolabs Japan) and subjected to the real-time qPCR using StepOnePlus Real-Time PCR system (Applied Biosystems) with KAPA SYBR FAST qPCR master mix (Kapa Biosystems). The qPCR conditions were as follows: initial holding at 95 °C for 20 s, 40 cycles of 95 °C for 3 s, 60 °C for 30 s, followed by 95 °C for 15 s, 60 °C for 1 min, 95 °C for 15 s. A melting curve was generated to confirm the specificity of the reactions. Each sample was tested in triplicates. The relative expression level of target genes was calculated by comparative CT (ΔΔCT) method using EF1α as the internal reference and a common standard sample (a mixture of RNAs from untreated SR1) for the normalization among assay plates. The primers used for qRT-PCR analysis are listed in Table S2.

### Determination of cell death

Cell death was measured using the Trypan Blue Assay as described previously (Morel and Dangl, 1999) with slight modifications. Three-week-old transgenic and control plants were Dex- or solvent-treated, kept for seven days, and observed for the phenotypic changes. The Agroinfiltration–mediated transient expression of *Tomato bushy stunt virus* (TBSV) P19 (Voinnet *et al*., 2003) were analyzed 2 days post-infiltration as a positive control for cell death (Scholthof *et al*., 1995). Leaf disks of 6 mm in diameter were heated for three minutes in trypan blue solution, cooled to room temperature, de-colorized using chloral hydrate solution, and mounted in 50% glycerol for microscopy analysis. For the quantitative determination of cell death, electrolyte leakage assay was conducted as described previously (Mackey *et al*., 2003) with slight modifications. Eight leaf disks from each plant were floated in 9 ml sterilized distilled water (SDW) and kept for 24 hours under dark conditions. The conductance of the leaked electrolyte in SDW was measured using a conductivity meter (LAQUAtwin-EC-11, HORIBA Scientific, Japan). After the measurement, leaf disks and SDW were recombined, boiled for 10 minutes, and the conductivity of total electrolyte was measured. The cell death was evaluated with the relative conductivity, the ratio of those for leaked to total electrolyte in three triplicate experiments. Statistical analyses [Tukey’s HSD (honestly significant difference) test] were performed using SPSS (Version 17) and Microsoft Office Excel 2016.

### DAB (3,3’-diaminobenzidine) staining for hydrogen peroxide detection

For *in situ* detection of hydrogen peroxide (H_2_O_2_), we performed 3,3’-diaminobenzidine (DAB) staining as described previously (Daudi and O’Brien, 2012) with slight modifications. Three-week-old transgenic and control plants were Dex- or solvent-treated and further grown for 24 hours. Leaf-disks of 6 mm in diameter were vacuum-infiltrated with 1 mg/mL DAB solution containing 0.05% Tween-20 in 50 mL conical tubes, kept on wet filter papers in Petri dishes, and incubated under light (70-100 μmolem^−2^s^−1^ for 30 minutes for test plants and 250-300 μmolem^−2^s^−1^ for 60 minutes for positive control plants), and for additional 3 hours 30 minutes in the dark. The leaf disks were de-colorized by boiling in 99.5% ethanol for 5-10 min and observed.

## Results

### RNA sequencing, mapping, and identification of DEGs

We previously reported that the silencing of HSP90C in transgenic tobacco lines resulted in chlorosis and growth suppression (Fig. 1A) accompanied by an activation of PR genes (Bhor *et al*., 2017b). RNA-seq analysis was conducted to elucidate the mechanisms underlying the development of chlorosis in this model system. Three-week-old i-hpHSP90C transgenic line H-4 and non-transformant SR1 were Dex- or control-treated, and RNA was extracted at 24 h post-Dex treatment from four individual plants from each plant/treatment group. Three each of them were selected for RNA-seq based on their integrity and purity (data not shown). The RNA sequencing gave 23 M reads/sample on average from the 12 samples, 80-90% of which were mapped to the tobacco reference transcriptome (Supplementary table S3).

**Fig. 1.**
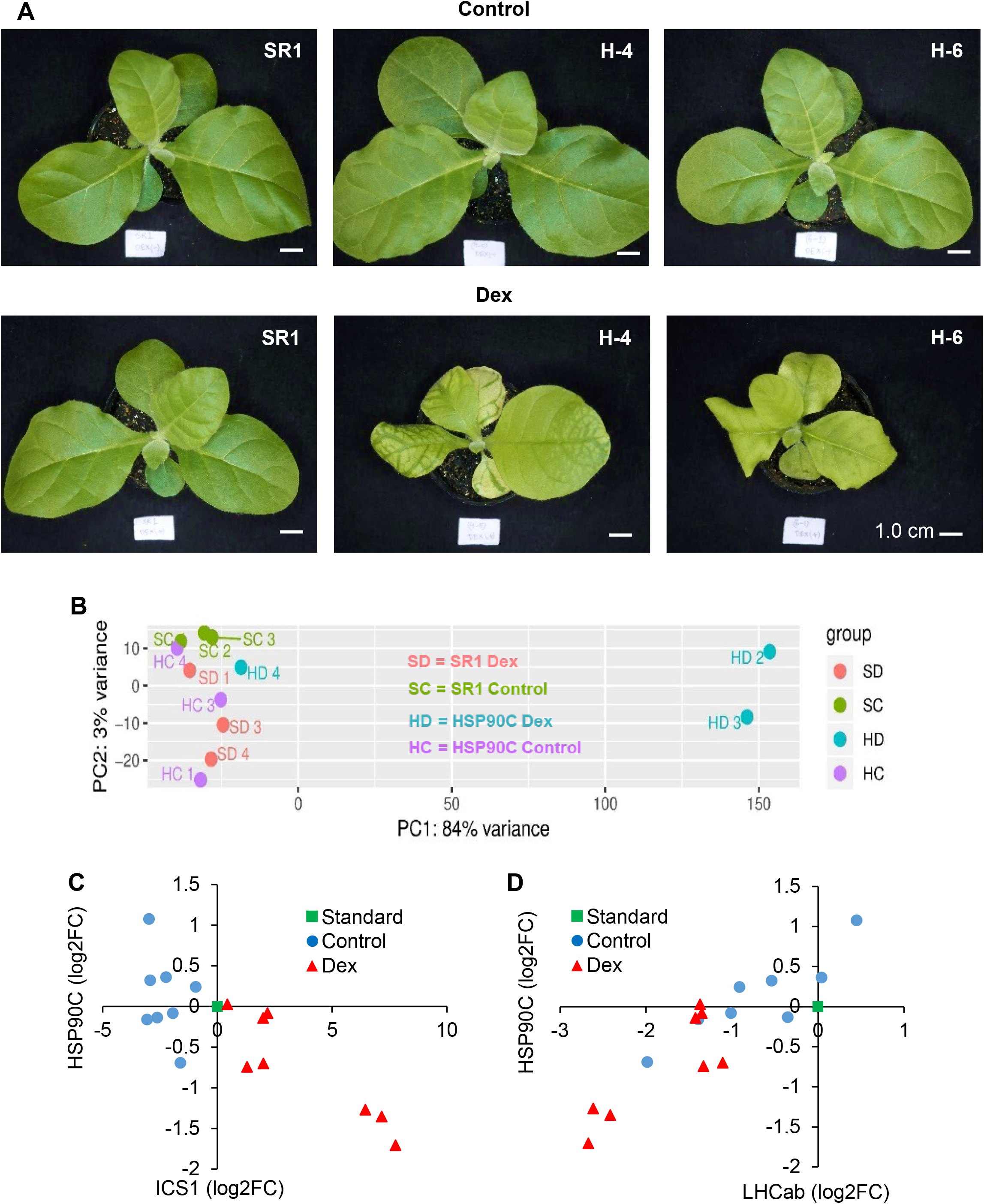
Phenotype and transcriptome change after induced silencing of HPS90 in transgenic tobacco plants. (A) Non-transformed SR1 (SR1), i-hpHSP90C-4 (H-4) and i-hpHSP90C-6 (H-6) plants were grown for three weeks, sprayed with 0.01% Tween-20 containing 0.5% ethanol (Control) or 50μM Dex (Dex), and photographed at 7 days post-treatment (dpt). Scale bars indicate 1 cm. (B) Principle Component Analysis of RNA-seq data. RNA preparations from 1-dpt control-treated SR1 (SC2, 3, and 4) and Dex-treated SR1 (SD1, 3, and 4), control-treated i-hpHSP90C-4 (HC1, 3, and 4) and Dex-treated i-hpHSP90C-4 (HD2, 3, and 4) were analyzed using DESeq2. (C and D) Variation of HSP90C silencing in individual Dex-treated i-hpHSP90C-4 plants. Expression levels of *ICS1* (C) and *LHCab* (D), representatives of up- and down-regulated genes found in the RNA-seq analysis, respectively, were plotted with those of *HSP90C*. The expression levels of these genes relative to a fixed standard sample (square) were quantified in eight individual plants each of control- (circles) and Dex-treated (triangles) i-hpHSP90C-4 plants.

In our initial analysis, the overall similarity within samples was evaluated by a principal component analysis (PCA) (Fig. 1B). The result showed that two samples, HD2 and HD3, showed clear differences from the control samples including the non-transformed SR1 samples regardless of the treatment and Dex-untreated i-hpHSP90C plants. In contrast, one of the Dex-treated i-hpHSP90C plants (HD4) showed the least differences from those control samples (Fig. 1B). The higher normalized HSP90C transcript counts indicated that the HD4 plant had not efficiently been silenced in HSP90C expression (Fig. 2D), suggesting that the transcriptomic changes are observed only after a significant HSP90C down-regulation. To test the hypothesis, we compared the expression levels by qRT-PCR of HSP90C and representative genes that show clear up- and down-regulation found in the RNA-seq analysis (see below), the isochorismate synthase 1 gene (*ICS1*) and a light-harvesting chlorophyll a/b-binding protein (*LHCab*), respectively. The gene expression of *ICS1* and *LHCab* correlated with the HSP90C expression negatively and positively, respectively (Fig. 1C and D). Importantly, in some plants of the i-hpHSP90C H-4 line, in which HSP90C expression levels were more than half of the control plants, the up-regulation of *ICS1* and the down-regulation of *LHCab* were not very prominent and within the variability of control (Dex-untreated) plants. In contrast to the H-4 line, the H-6 line of i-hpHSP90C transgenic plants showed highly reproducible Hsp90C silencing and the up- and down-regulation of *ICS1* and *LHCab*, respectively (Supplementary Fig. S1). The SR1 plant did not respond to Dex-treatment in the expression of the genes examined. These results support the hypothesis above, and therefore, we omitted the HD4 sample from the differential expression analysis.

**Fig. 2.**
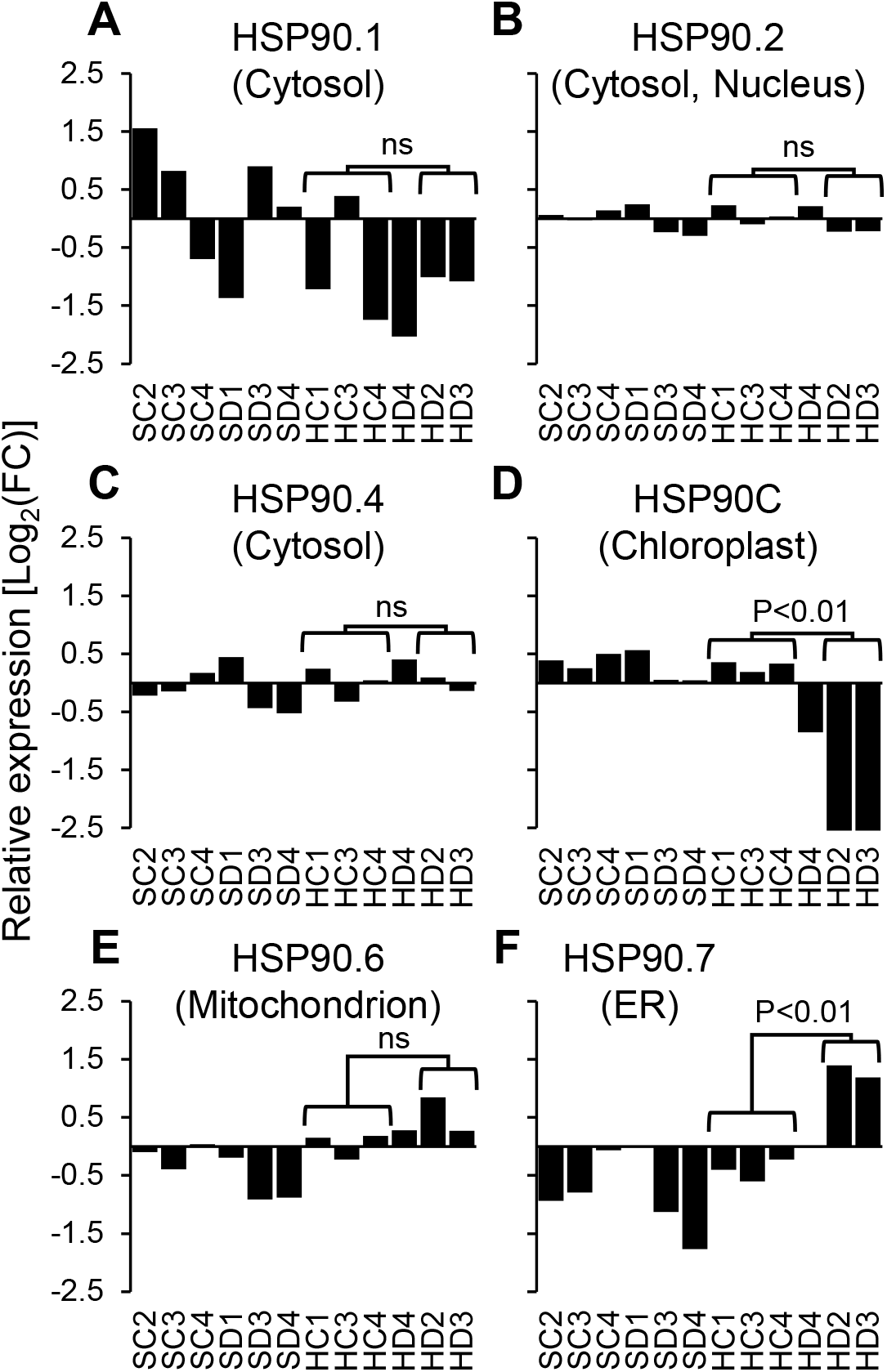
Expression changes of HSP90 family genes in RNA-seq samples. Relative normalized read counts in all RNA-seq samples are shown for tobacco homologs of Arabidopsis HSP90 family genes: cytosolic HSP90.1 (A), nucleo-cytosolic HSP90.2 (B), cytosolic HSP90.4 (C), chloroplast HSP90C or HSP90.5 (D), mitochondrial HSP90.6 (E), ER-localized HSP90.7 (F). Statistical significance of the difference in read counts was evaluated between control-treated (HC1, 3, and 4) and Dex-treated (HD2, and 3) in DESeq2 analysis. ns, not significant. Data from the HD4 sample was omitted as mentioned in the text.

The gene expression of HD2 and HD3 samples (HD) were compared using DESeq2 with all three groups, untreated line H-4 (HC), Dex-treated SR1 (SD), and untreated SR1 (SC), which have never shown chlorosis in our repeated experiments. DEGs were picked up from the DESeq2 data based on the combined criteria of log2(FC) values below −1 or above 1, and the adjusted p-values lesser than 0.05. The MA plots support that three comparisons identified a consistent set of DEGs (Supplementary Fig. S2). The analyses of differentially expressed mRNA transcripts in three different comparisons above—HD vs HC, HD vs SD, and HD vs SC—identified 7267 (55.9%) and 8042 (47.4%) commonly up- and down-regulated mRNAs, respectively (Fig. 3A and B). Because the mRNA IDs in the tobacco reference transcriptome are not readily used for downstream GO enrichment analysis, those annotated with the *Arabidopsis* AGI codes were selected. Out of the differentially expressed mRNAs above, 4896 up-regulated mRNAs had annotation with 2746 different AGI codes comprising 61.4% of DEGs with AGI annotation common in three comparisons, and 6307 down-regulated mRNAs had 3490 (59.0%) AGI codes (Fig. 3C and D and Supplementary tables S4 & S5).

**Fig. 3.**
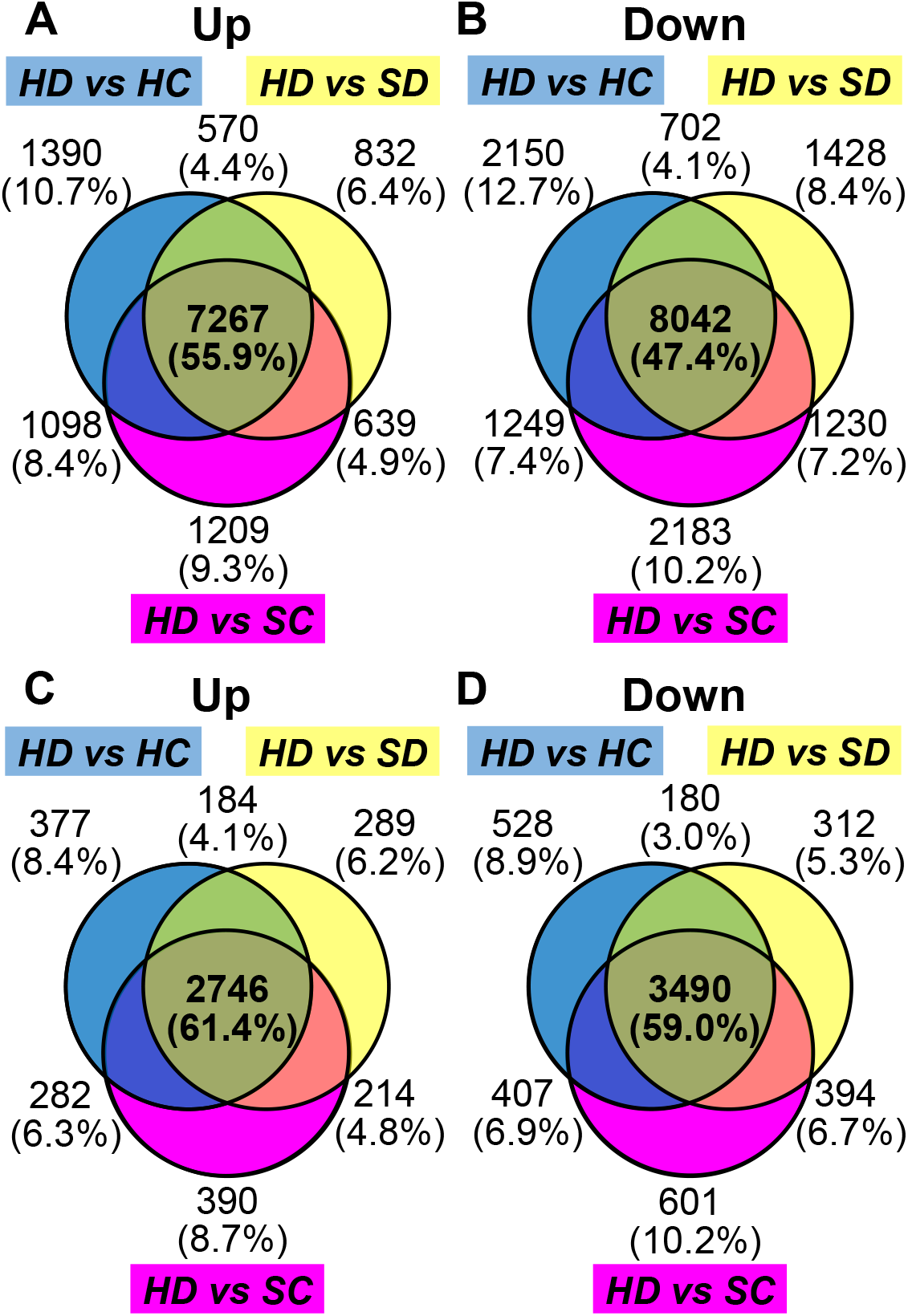
Venn diagrams of differentially expressed genes. Results from differential expression analysis with DESeq2 were made in three different comparisons: Dex-treated H-4 vs control H-4 (HD vs. HC), Dex-treated H-4 vs Dex-treated SR1 (HD vs. SD), and Dex-treated H-4 vs Control SR1 (HD vs. SC). The differentially expressed genes (DEGs) were detected using DESeq2 as mentioned in Materials and methods. A and B show DEGs detected in the analysis with mRNA ID in the reference transcriptome, and C and D show those annotated with AGI codes. A and C show numbers of upregulated genes whereas B and D show those of downregulated genes.

### Functional classification of DEGs

The AGI codes given to the up- and down-regulated mRNAs were used for the GO enrichment analysis. Tables 1 and 2 present the selected GO terms for biological processes enriched in the lists of up- and down-regulated mRNAs, respectively. The enriched GO terms in the up-regulated genes include innate immune response, response to wounding, response to oxidative stress, response to phytohormones [salicylic acid (SA), jasmonic acid (JA), and abscisic acid(ABA)], and hypersensitive cell death (Table 1 & Supplementary table S6). In accordance with the observation, biosynthetic genes of SA and JA were found upregulated (Table 1). The heatmap (Fig. 4) shows the expression changes of genes selected from those in GO terms, response to SA (GO:0009751), response to JA (GO:0009753), response to oxidative stress (GO:0006979) and cell death (GO:0008219), which are indicated by double-headed arrows on the right. The results suggest that the reduced supply of HSP90C to chloroplast elicits defense response involving some phytohormone pathways.

**Table 1.** Selected GO terms (Biological process) for upregulated genes in i-hpHSP90C transgenic line after Dex treatment.

| **GO Terms** | **Total** | **DEGs** | **Fold enrich.** | **FDR** |
| --- | --- | --- | --- | --- |
| **Immune system process** | 372 | 101 | 2.73 | <0.01 |
| Innate immune response | 300 | 83 | 2.78 | <0.01 |
| **Response to stimulus** | 5327 | 880 | 1.66 | <0.01 |
| Response to stress | 3079 | 547 | 1.79 | <0.01 |
| Response to osmotic stress | 545 | 105 | 1.94 | <0.01 |
| Response to salt stress | 469 | 90 | 1.93 | <0.01 |
| Defense response | 1005 | 231 | 2.31 | <0.01 |
| Response to wounding | 207 | 59 | 2.86 | <0.01 |
| Response to oxidative stress | 392 | 83 | 2.13 | <0.01 |
| Response to hormone | 1360 | 235 | 1.74 | <0.01 |
| Response to SA | 206 | 41 | 2 | <0.01 |
| Response to JA | 217 | 46 | 2.13 | <0.01 |
| JA mediated signaling pathway | 67 | 23 | 3.45 | <0.01 |
| Response to ABA | 524 | 103 | 1.98 | <0.01 |
| Response to ER stress | 97 | 37 | 3.83 | <0.01 |
| ERAD pathway | 60 | 17 | 2.85 | 0.01 |
| ER unfolded protein response | 36 | 13 | 3.63 | 0.01 |
| **Metabolic process** | 7731 | 992 | 1.29 | <0.01 |
| Cellular respiration | 80 | 21 | 2.64 | 0.01 |
| Protein glycosylation | 104 | 24 | 2.32 | 0.01 |
| Protein autophosphorylation | 183 | 46 | 2.53 | <0.01 |
| JA metabolic process | 50 | 14 | 2.81 | 0.03 |
| JA biosynthetic process | 26 | 10 | 3.87 | 0.02 |
| Regulation of SA metabolic process | 20 | 10 | 5.02 | 0.01 |
| Regulation of SA biosynthetic process | 12 | 8 | 6.7 | 0.01 |
| **Cellular process** | 10043 | 1299 | 1.3 | <0.01 |
| Cell death | 112 | 32 | 2.87 | <0.01 |
| Programmed cell death | 91 | 27 | 2.98 | <0.01 |
| Plant-type hypersensitive response | 57 | 25 | 4.41 | <0.01 |
| Regulation of cell death | 79 | 22 | 2.8 | <0.01 |
| Negative regulation of cell death | 33 | 13 | 3.96 | <0.01 |
| **Plant organ development** | 946 | 141 | 1.5 | <0.01 |
| Root morphogenesis | 253 | 43 | 1.71 | 0.04 |
| Leaf senescence | 103 | 29 | 2.83 | <0.01 |

**Table 2.** Selected GO terms (Biological Process) for downregulated genes in i-hpHSP90C transgenic line after Dex treatment.

| GO Terms | Total | DEGs | Fold enrich. | FDR |
| --- | --- | --- | --- | --- |
| **Metabolic process** | 7731 | 1278 | 1.3 | <0.01 |
| Carbohydrate metabolic process | 668 | 146 | 1.72 | <0.01 |
| Lipid metabolic process | 728 | 175 | 1.89 | <0.01 |
| Cellular AA* metabolic process | 333 | 85 | 2.01 | <0.01 |
| Photosynthesis | 177 | 102 | 4.54 | <0.01 |
| Cofactor metabolic process | 416 | 129 | 2.44 | <0.01 |
| Pigment metabolic process | 123 | 48 | 3.07 | <0.01 |
| Vitamin metabolic process | 77 | 31 | 3.17 | <0.01 |
| **Cellular process** | 10043 | 1592 | 1.25 | <0.01 |
| Cell wall organization | 279 | 60 | 1.69 | 0.01 |
| Cell cycle | 486 | 106 | 1.72 | <0.01 |
| Cellular homeostasis | 311 | 62 | 1.57 | 0.04 |
| Cellular component organization | 2503 | 454 | 1.43 | <0.01 |
| Plastid organization | 298 | 136 | 3.6 | <0.01 |
| **Response to stimulus** | 5327 | 903 | 1.34 | <0.01 |
| Response to osmotic stress | 545 | 103 | 1.49 | 0.01 |
| Response to oxidative stress | 392 | 87 | 1.75 | <0.01 |
| Response to salt stress | 469 | 91 | 1.53 | 0.01 |
| Response to auxin | 310 | 72 | 1.83 | <0.01 |
| Response to light stimulus | 680 | 198 | 2.29 | <0.01 |
| **Rhythmic process** | 122 | 43 | 2.78 | <0.01 |
| Circadian rhythm | 109 | 42 | 3.04 | <0.01 |

\* AA = Amino Acid

**Fig. 4.**
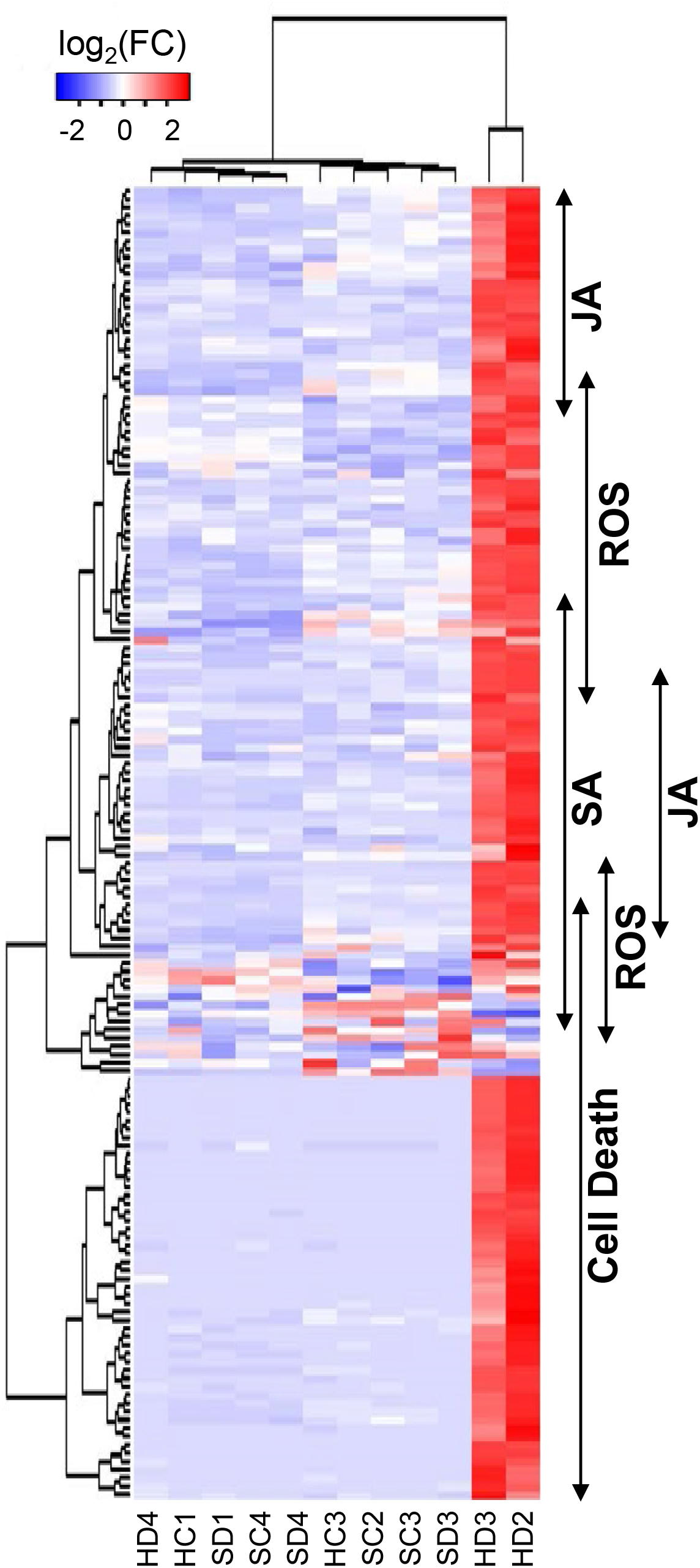
Hierarchical clustering of upregulated DEGs with selected annotations. The genes annotated with GO terms, response to SA (GO:0009751), response to JA (GO:0009753), response to oxidative stress (GO:0006979), and cell death (GO:0008219) were selected (Supplementary Table S9), and their relative-normalized count data were used to draw a heatmap. Each column represents a sample, and each row represents a gene selected. Differences in the expression are shown in different colors, red and blue represent the up- and down-regulated expression, respectively.

In addition to the activation of defense response, genes involved in the response to endoplasmic reticulum (ER) stress were remarkably upregulated after the impaired supply of HSP90C (Table 1). This observation could be attributed to the non-specific silencing of ER-localizing HSP90 family protein by the HSP90C hpRNA. Therefore, we examined the expression levels of six out of seven HSP90 family proteins in the RNA-seq data because no tobacco mRNA in the reference transcriptome was annotated with AT5G56010 encoding a nucleo-cytosolic HSP90.3. The true target of hpRNA-mediated silencing, HSP90C or HSP90.5, was significantly downregulated as shown (Fig. 2D), but the expression of cytosolic HSP90.1 and HSP90.4, nucleo-cytosolic HSP90.2, and mitochondrial HSP90.6 was not affected by the Dex-treatment (Fig. 2A, B, C, and E). In contrast, statistically significant upregulation in Dex-treated HD2 and HD3 was detected in ER-localizing HSP90.7 (Fig. 2F). The results contradict the off-target silencing of ER-localizing HSP90, and thus, suggest the interplay between the protein quality control system of ER with that in plastid or immune responses.

The enriched GO terms (Biological process) in the downregulated genes include photosynthesis, pigment metabolic process, and plastid organization (Table 2 & Supplementary table S7), which is consistent with crucial roles of HS90C in chloroplast biogenesis. In addition, some primary metabolism genes annotated with GO terms, carbohydrate metabolic process, lipid metabolic process and cellular amino acid metabolic process, other metabolic genes annotated with cofactor metabolic process and vitamin metabolic process, and some cellular process genes annotated with cell cycle, cell wall organization, and cellular homeostasis were shown to be downregulated (Table 2 & Supplementary table S7), which is consistent with the upregulation of cell death-related genes (Table 1 & Supplementary table S6). Interestingly, three GO terms, response to osmotic stress, response to oxidative stress, and response to salt stress were enriched in the downregulated genes. However, they were also enriched in the upregulated genes (Tables 1 and 2). The results suggest that impaired HSP90C supply would induce drastic switching in these stress-responsive genes (Supplementary Fig. S3).

### Detection of cell death and reactive oxygen species in HPS90C-silenced plants

The significant upregulation of the cell death pathway prompted us to detect cell death in i-hpHSP90C lines. Because lower (older) leaves showed severer chlorosis than the upper (younger) leaves, we examined the cell death in those leaves separately. The trypan blue staining showed light blue stain in upper leaves and more intense staining in older leaves, albeit to a lesser extent than the positive control, of Dex-treated H-4 and H-6 plants (Fig. 5J–L and P–R). In the positive control plants, in which TBSV P19 had transiently been expressed for two days, some leaf discs were cut out to include both dead and living parts to make the difference in staining of those parts clear (Fig. 5S and T). The staining of lower leaves from Dex-treated H-4 and H-6 plants was not uniform, suggesting the cell death induction in those leaves were sporadic within the leaf tissue. Electrolyte leakage assay confirmed the cell death in lower leaves but not that in upper leaves (Fig. 5U). Microscopic observation of trypan blue stained leaf tissue indicated that dead cells in lower leaves of Dex-treated H-4 and H-6 plants were barely shrunken (Fig. 5h and l), unlike those in the positive control (Figure 5m). The upper leaves exhibited small patches of dead cells, suggesting the sporadic and age-dependent natures of cell death in Dex-treated H-4 and H-6 plants (Fig. 5g and k). A small fraction of cells in lower but not upper leaves of untreated H-6 showed blue staining (Fig. 5j), suggesting that the hpRNA to *HSP90C* had been expressed in a leaky fashion in a small number of cells in untreated H-6 line and that the cell death observed is age-dependent. Dex-treated and untreated control plants (SR1) and untreated H-4 did not show any sign of cell death (Fig. 5a–d, e, and f). These results collectively suggest that the death of cells, in which HSP90C supply is impaired, is stochastically initiated and proceeds somehow slowly, although the RNA-seq suggests that it is a plant-type hypersensitive response (Table 1).

**Fig. 5.**
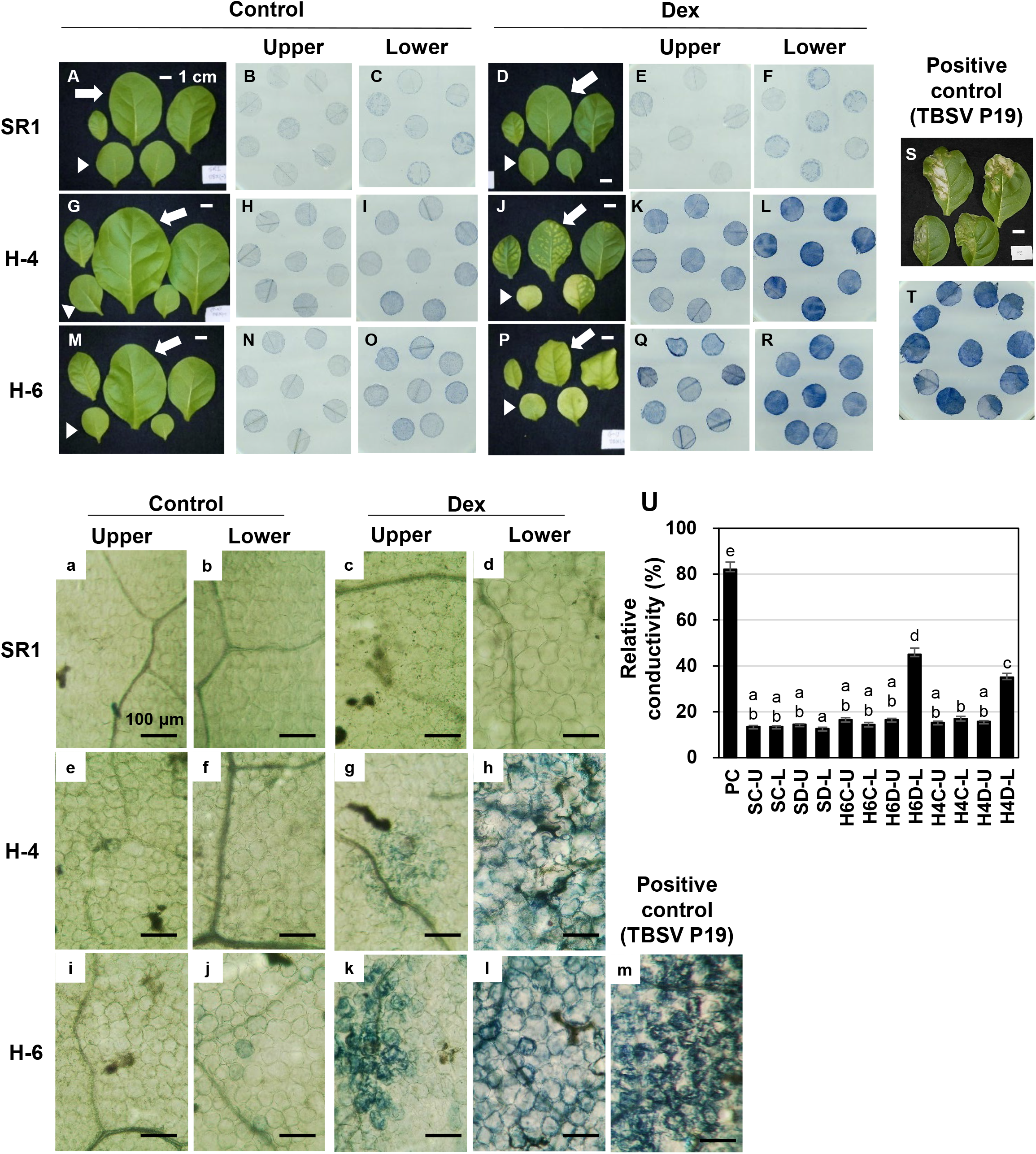
Detection of cell death in chlorotic leaves. Control plants (SR1; A–F, and a–d), i-hpHSP90C transgenic line 4 (H–4; G–L and e–h), and line 6 (H–6; MR and i–l) plants were grown, treated with control or Dex solution. They were harvested and photographed at 7 dpt (A, D G, J, M, P, and S; representative plants from triplicate experiments; scale bars denote 1 cm) followed by cell death assays. SR1 leaves transiently expressing TBSV P19 for 2 days served as a positive control (S, T, and m). Cell death assay was performed in upper leaves (pointed by arrows) and lower leaves (pointed by arrowheads) Leaf disks of 6 mm in diameter were stained with trypan blue and photographed at macroscopic (B, C, E, F, H, I, K, L, N, O, Q, R, and T) and microscopic (a–m; scale bars denote 100 μm) levels. U, quantification of cell death by an electrolyte leakage assay. PC, positive control with transient TBSV P19 expression; SC, control-treated SR1; SD, Dex-treated SR1; H4C, control-treated line 4; H4D, Dex-treated line 4; H6C, control-treated line 6; H6D, Dex-treated line 6; U, upper leaves; L, lower leaves. Error bars denote standard deviations in triplicate experiments. Different letters indicate statistically significant difference between treatments (LSD test, P<0.05). The experiment was repeated at least three times.

The upregulation of genes involved in the response to oxidative stress (Table 1 and Fig. 4) suggests the production of reactive oxygen species (ROS) in Dex-treated H-4 and H-6 plants. Because ROS is well studied as a signal mediator and an executor of plant cell death, we tried to detect H_2_O_2_ as a representative of ROS. Positive control with intense light resulted in the accumulation of brownish DAB precipitate in chloroplasts (Fig. 6M). Intense DAB staining of chloroplasts was barely visible in Dex-treated and untreated control plants (SR1), and untreated H-4 and H-6 (Fig. 6A–D, E, F, I and J). Evident DAB staining of chloroplasts was observed in leaves of Dex-treated H-4 and H-6 plants (Fig. 6G, H, K, and L). The results suggest that, upon the loss of sufficient levels of HSP90C, chloroplasts produce ROS, which would trigger the cell death response. Although the overall DAB pigmentation was more intense in lower leaves, especially in H-4, pigmentation of each chloroplast was comparable between upper and lower leaves. The observation suggests that the extents of cell death correlate with the number of cells producing ROS but not with the magnitude of ROS production.

**Fig. 6.**
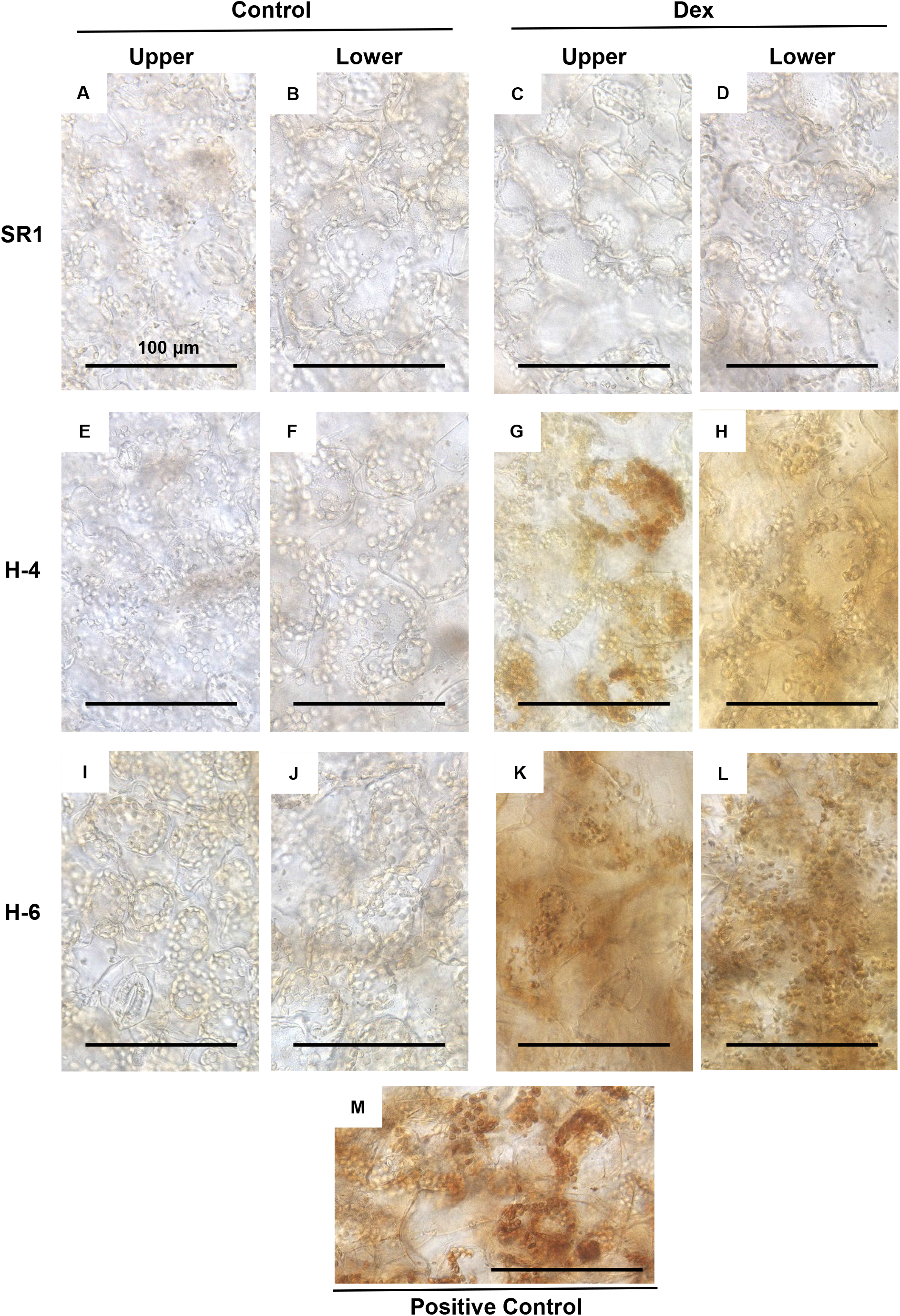
H_2_O_2_ production in the chloroplasts of leaves developing chlorosis. Control plants (SR1; A–D), i-hpHSP90C transgenic line 4 (H–4; E–H), and line 6 (H–6; I–L) plants were grown, treated with control or Dex solution. At 24 hrs post-treatment, leaf disks of 6 mm in diameter were cut out from upper and lower leaves, vacuum-infiltrated with DAB solution, and incubated under light conditions (70-100 μmolem^−2^s^−1^ for 30 minutes) Untreated SR1 plants illuminated with 250-300 μmolem^−2^s^−1^ for 60 minutes served as a positive control. All leaf disks were incubated under dark for 3.5 hrs, de-colorized, and observed under a microscope.

## Discussion

We previously established an inducible silencing system for HSP90C in tobacco and confirmed that the silencing of HSP90C alone could lead plants to chlorosis (Bhor *et al*., 2017*b*). The system would have an advantage over the experimental systems with the virus- or viroid-infected plants in the analysis of mechanisms underlying the development of disease symptom-like phenotype such as chlorosis. It is possible in this system to analyze plant cells that have committed to developing but not exhibited chlorosis. Exploiting the advantage in the present study, we explored the early molecular changes leading to the development of chlorosis using RNA-seq analysis. Although studies have shown the transcriptome changes in virus- and viroid-infected plants (Mochizuki *et al*., 2014; Xia *et al*., 2017; Zheng *et al*., 2017), the strength of the present study is that we could detect any changes that precede detectable chlorosis.

From two independent transgenic lines, H-4 and H-6, which show mild and severe chlorosis and growth suppression after Dex treatment, respectively, we selected the former for the RNA-seq analysis. In the triplicate experiment, an individual Dex-treated of H-4 plant (HD4 in Fig. 1B and Fig. 2) showed a moderate reduction in HSP90C expression levels (Fig. 2D) and the lack of transcriptome changes observed in other two Dex-treated H4 plants (Fig. 1B, Fig. 4, and supplementary Fig. S3). Because such a variation within individual Dex-treated H-4 plants was confirmed by qRT-PCR (Fig. 1C and D), we omitted the HD4 data from our RNA-seq data analysis. Although more biological replicates are recommended for reliable RNA-seq data analysis in general, DEseq2 used in this study has been shown to give the least false positive rate (Schurch *et al*., 2016) Therefore, most of the data analysis results were taken into account in this report. In contrast to Dex-treated H-4 plants, Dex-treated H-6 plants showed drastic downregulation of *LHCab* expression and dramatic induction of *ICS1* expression. The results suggest a more rapid progression of molecular changes toward chlorosis in H-6 than in H-4 plants, which may be more suitable than H-6 for analyzing the early molecular changes.

We found several characteristic transcriptome changes in HD2 and HD3 plants, which we believed to be on the way to develop chlorosis. Firstly, the downregulation was observed in genes involved in (group 1) photosynthesis and plastid organization, (group 2) primary and secondary metabolisms, and (group 3) cell and plant growth. Secondly, the genes involved in (group 4) immune response accompanying cell death, and (group 5) ER stress response. Finally, in the gene involved in response to different abiotic stresses, a subset was upregulated, and the other was downregulated.

Among the transcriptome changes above, downregulation of chloroplast and photosynthesis-related genes (CPRGs) or group 1 genes has been widely reported in symptomatic tissues of different combinations of host plants and viruses (Satoh *et al*., 2010; Postnikova and Nemchinov, 2012; Mochizuki *et al*., 2014; Zanardo *et al*., 2019). Although it is natural that the expression of CPRGs is downregulated in chlorotic tissue, the present study, together with our previous results (Waliullah *et al*., 2014, 2015; Bhor *et al*., 2017*a*,*b*) strongly suggest that the CPRGs downregulation preceding visible chlorosis is the primary pathway of chlorosis development. The mechanism underlying the CPRGs downregulation may differ within pathosystems, but the present study suggests a possible involvement of retrograde signaling (RS) in chlorosis development induced by sub-viral pathogens. The downregulation of groups 2 and 3 genes would also be attributed to the RS activation, which reprograms transcriptome from growth and differentiation state to stress response state (Crawford *et al*., 2018). The RNA-seq data suggest the activation of RS pathways as manifested by the increased expression of transcription factors involved in RS-mediated transcriptome changes and decreased expression of those involved in chloroplast biogenesis (Supplementary Table S8) (Leister *et al*., 2014). Given RS pathways are activated in the chlorosis model of the present study, ROS could be a primary signal as we detected H_2_O_2_ production in chloroplasts in a day after the induction of HSP90C silencing. H_2_O_2_ molecules are assumed to move across the biological membrane (Henzler and Steudle, 2000), and thus, would move from chloroplast to nucleo-cytoplasmic space to activate different signaling pathways. Although transcriptome profile and ROS detection support the RS activation during chlorosis development, other RS signaling molecules should be analyzed to confirm the activation of RS pathways in the present model system. It is noteworthy that cyanobacterial HSP90, HtpG, interacts with and modulates the activity of uroporphyrinogen decarboxylase and thus regulating tetrapyrrole biosynthesis (Saito *et al*., 2008).

Among group 4 or immunity genes, *ICS1* was highly upregulated in the chlorosis model (Fig. 1C, Supplementary Fig. S1, Supplementary Tables S4, and S6). In addition to pathogen attack and UV irradiation, *ICS1* expression is reportedly upregulated with excess light and β-cyclocitral, a retrograde signaling molecule produced by the oxidation of β-carotene with singlet oxygen (Lv *et al*., 2015). The upregulation of *ICS1* expression in the present model system could also be regulated by some signal(s) from chloroplast to the nucleus. Our RNA-seq data has indicated significant upregulation of three transcription factors, SARD1, CBP60g, WRKY28, which have significant roles in ICS1 gene activation (Zhang *et al*., 2010; van Verk *et al*., 2011; Wang *et al*., 2011; Sun *et al*., 2015) and their flg22-induced expression reportedly depends on chloroplast-localized calcium sensor, CAS (Nomura *et al*., 2012). Because cytoplasmic calcium-dependent protein kinases have shown to regulate those transcription factors (Boudsocq and Sheen, 2013; Poovaiah *et al*., 2013), the importance of chloroplast signal in ICS1 induction needs to be clarified by further study.

ICS1 is the key enzyme of SA biosynthesis, which is required for both local and systemic acquired resistance, while SA synthesized through this pathway seems to potentiate plant cell death (Nawrath and Métraux, 1999; Gross *et al*., 2006; Strawn *et al*., 2007; Garcion *et al*., 2008; Chen *et al*., 2009). It is well known that SA induces cell death in plants as a defense response (Brodersen *et al*., 2005; Radojičić *et al*., 2018). We observed the H_2_O_2_ production in the chloroplast of Dex-treated i-hpHSP90C plants (Fig. 5) and the upregulation of ROS-responsive genes (Fig. 4, Supplementary Table S6). ROS produced in the chloroplast (Fig. 6) can induce SA biosynthesis most likely through ICS1 gene activation (Wildermuth *et al*., 2001; Garcion *et al*., 2008; Herrera-Vásquez *et al*., 2015). Taken together, the cell death we observed in the chlorosis model plants (Fig. 5) is suggested to be first triggered by the ROS production and then activated through SA-ROS self-amplification loop (Vlot *et al*., 2009). Although with the least plant growth, our preliminary experiment of plant culture in the dark suggests the chlorosis development requires the light-dependent ROS production (Supplementary Fig. S4). Studies have reported that light-dependent ROS production in chloroplast leads to the HR-like cell death and antiviral resistance (Chandra-Shekara *et al*., 2006; Liu *et al*., 2007; Chen *et al*., 2015; Hamel *et al*., 2016). However, the significance of SA in cell death in the present model system needs to be studied further.

In addition to defense genes, we found that ER stress response genes were upregulated in the chlorosis model (Table 1). Although physical interaction between chloroplasts and ER has been demonstrated, no protein transport was confirmed between these organelles (Schattat *et al*., 2012; Barton *et al*., 2018; Liu and Li, 2019). Therefore, it is unlikely that loss of protein quality control in chloroplast could directly induce the unfolded protein response (UPR). The indirect induction of UPR by the impaired HPS90C supply would involve either retrograde signaling or stress hormone response. A retrograde signal molecule, methylerythritol cyclodiphosphate (MEcPP), has been shown to induce UPR in ER (Walley *et al*., 2015; Benn *et al*., 2016) and SA has been shown to activate the major IRE1-bZIP60 pathway of UPR (Nagashima *et al*., 2014; Park and Park, 2019). Further study of the mode of induction of UPR would provide us with insight into its role in chlorosis development.

We summarize the molecular events discussed above that lead to the development of chlorosis after the induced silencing of the *HSP90C* gene (Fig. 7). Reduced HSP90C supply would impair protein quality control in the chloroplast, hamper normal chloroplast function, and thus, lead to the production of ROS in the chloroplast. ROS would induce SA production, and they would develop a self-amplifying loop. ROS would also activate the chloroplast retrograde signaling, which results in the upregulation of stress-responsive genes, including pathogenesis-related (PR) genes, and the downregulation of CPRGs in the nucleus, which is most likely to be the primary cause of chlorosis. In addition, the HR-like cell death pathway would be activated by ROS, SA pathway, other stress hormone pathways, and/or UPR. It is well known that chlorosis induced by different cue ends up to the death of plant tissue. Therefore, the activation of the cell death response could have a role in chlorosis induction or acceleration of the processes leading to visible chlorosis. Although the importance of cell death induction in chlorosis remains to established, the present study sheds light on the importance of prior activation of the cell death pathway in chlorosis development.

**Fig. 7.**
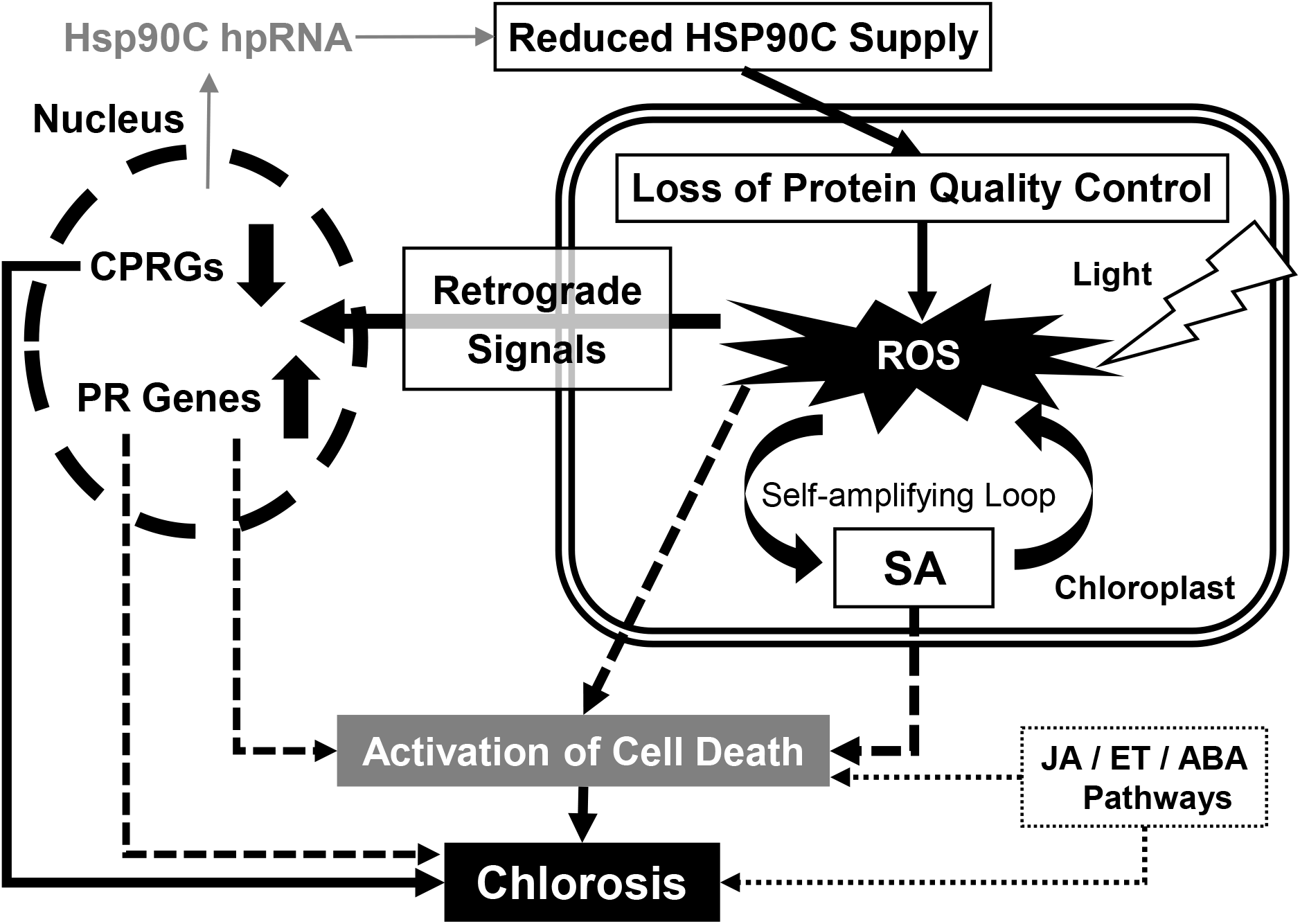
A schematic model for the molecular mechanism of chlorosis. Inducible silencing of the HSP90C gene would cause a reduction of HSP90C levels in the chloroplast, which would cause the loss of protein quality control. The loss of protein quality control would lead chloroplasts to produce ROS in a light-dependent manner. The effects of ROS and SA would be enhanced by a self-amplifying loop. ROS would serve as the chloroplast retrograde signal and activate other retrograde signaling pathways, leading to the upregulation of pathogenesis-related (PR) genes and the downregulation of CPRGs. ROS, SA, the activation of PR genes, and other stress responses including JA/ET/ABA pathways could stimulate the cell death pathway. The CPRGs downregulation would be the primary cause of chlorosis, and ROS-mediated triggering of HR-like cell death in the chlorotic tissues could have a role in the development of chlorosis. Solid line arrows indicate the steps with experimental supports, whereas broken line arrows are hypothetical.

In the case of a virus or viroid infection, the activation of the immune pathway must be unfavorable to the pathogens. Therefore, it is an acceptable idea that pathogens have some way to inhibit the defense response activated by the chloroplast stress. Indeed, PLMVd and some other viruses such as CMV induce clear bleaching-type chlorosis, in which cell death is unlikely (Mochizuki and Ohki, 2011; Navarro *et al*., 2012). It is notable that PLMVd infection reportedly produces siRNAs to several other host genes, including NB-LRR type disease resistance genes, in addition to those to HSP90C (Navarro *et al*., 2012; Chiumenti *et al*., 2018). The analysis of pathogens’ anti-defense strategies would pave the way for a better understanding of mechanisms underlying the development of virus disease symptoms.

## Acknowledgments

This study was supported in part by The United Graduate School of Agricultural Sciences, Ehime University, and JSPS KAKENHI grants 26292026, 15K14664, and 19K06055 to KK. RNA-seq analysis was conducted under the support of the Cooperative Research Grant of the Genome Research for BioResource, NODAI Genome Research Center, Tokyo University of Agriculture. Islam Shaikhul was supported by the MEXT.

## Abbreviations

CPRGs: Chloroplast and photosynthesis-related genes
DEGs: Differentially expressed genes
Dex: Dexamethasone
hp-RNA: Hairpin RNA
HSP90C: Chloroplast heat shock protein 90
ROS: Reactive oxygen species
siRNA: Small interfering RNA

## Supplementary Data

**Supplementary Fig. S1.** Comparison between the expression of HSP90C gene and selected DEGs in non-transformed control SR1 and i-hpHSP90C line 6 plants. X-axis shows the log2(FC) value (from qRT-PCR analysis) of two selected genes, ICS1 and LHCab, and Y-axis indicates the log2(FC) value of HSP90C gene relative to a common standard sample.

**Supplementary Fig. S2. (A-C)** MA plot showing the overview of the DEGs in two-group of comparison (i.e. Dex treated and untreated).

**Supplementary Fig. S3.** Heat maps showing the oxidative, osmotic, and salt stress-responsive genes. Each column represents a sample, and each row represents a gene selected. Differences in expression are shown in different colors, red and blue represent the up- and down-regulated expression, respectively.

**Supplementary Fig. S4.** Lack of chlorosis response in dark-grown i-hpHSP0C (6-1) plants.

**Supplementary Table 1**. Transcript ID and Gene ID mapping file.

**Supplementary Table 2.** Primers used for qRT-PCR analysis in the present study.

**Supplementary Table 3.** Summary of RNA-Seq data and mapping of the clean reads to *N. tabacum* TN90 reference transcriptome. S, SR1 (or non-transformant); H, i-hpHSP90C; D, Dex-treated; C, Control.

**Supplementary Table 4.** List of upregulated DEGs in chlorosis model plants (commonly upregulated DEGs in all three comparisons).

**Supplementary Table 5.** List of downregulated DEGs in chlorosis model plants (commonly downregulated DEGs in all three comparisons).

**Supplementary Table 6.** List of GO (Biological process) enriched with upregulated DEGs of chlorosis model plants.

**Supplementary Table 7.** List of GO (Biological process) enriched with downregulated DEGs of chlorosis model plants.

**Supplementary Table 8.** Expression levels of RS-related transcription factor genes.

