## Supplemental Data 1 for "Impaired expression of chloroplast HSP90C chaperone activates plant defense responses leading to a disease symptom-like phenotype"

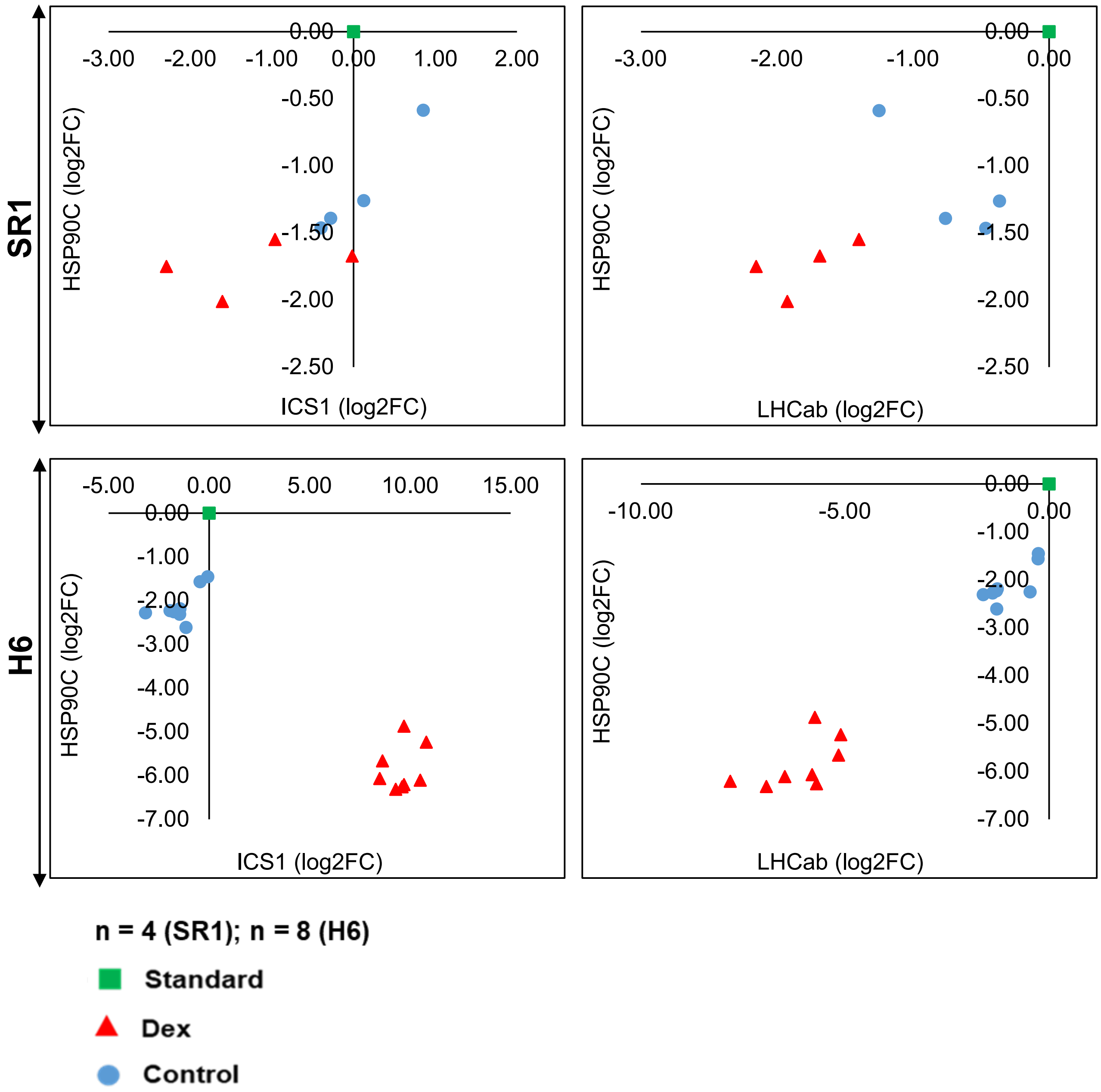


Supplementary Figure 1

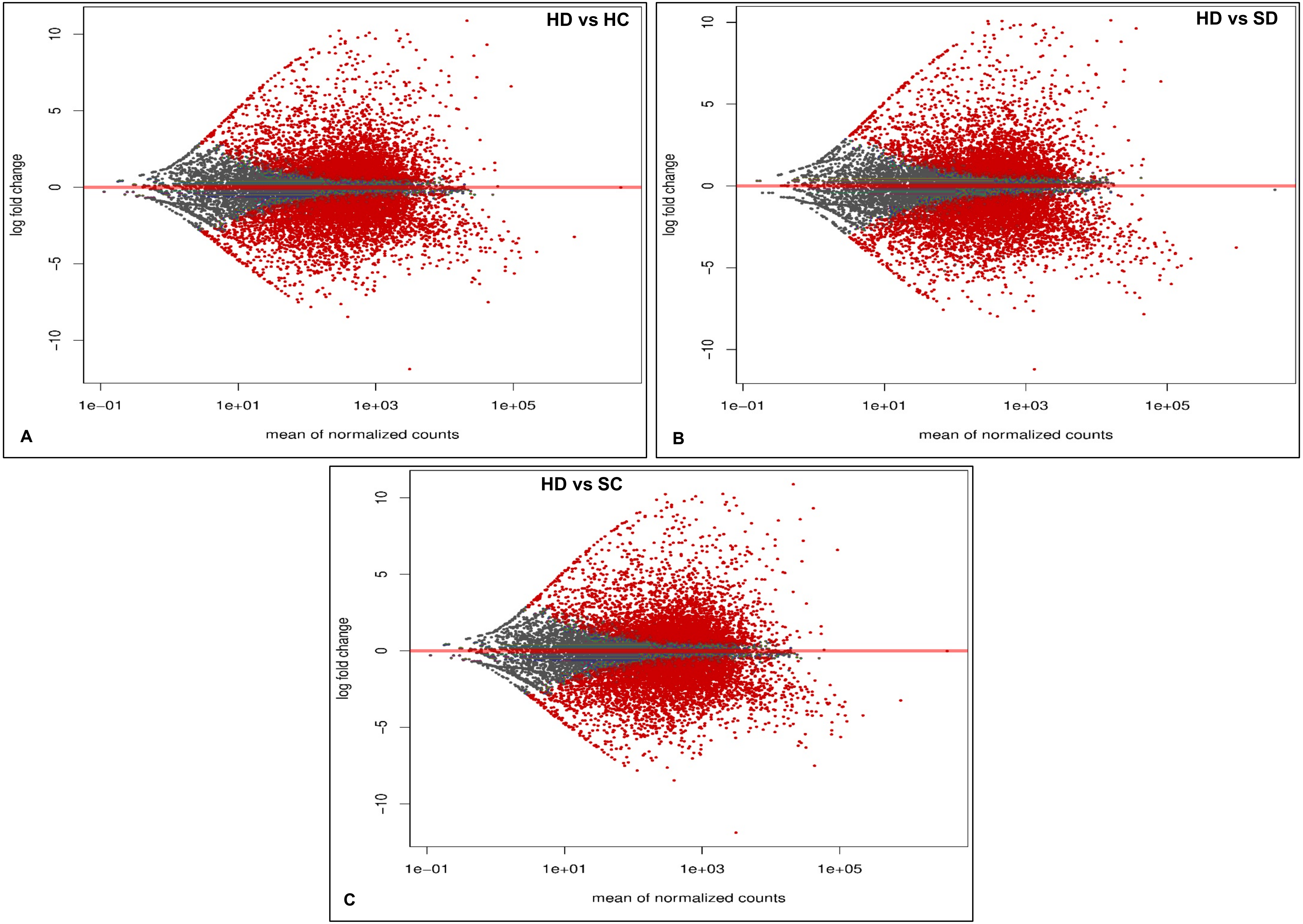


**Supplementary Figure 2**

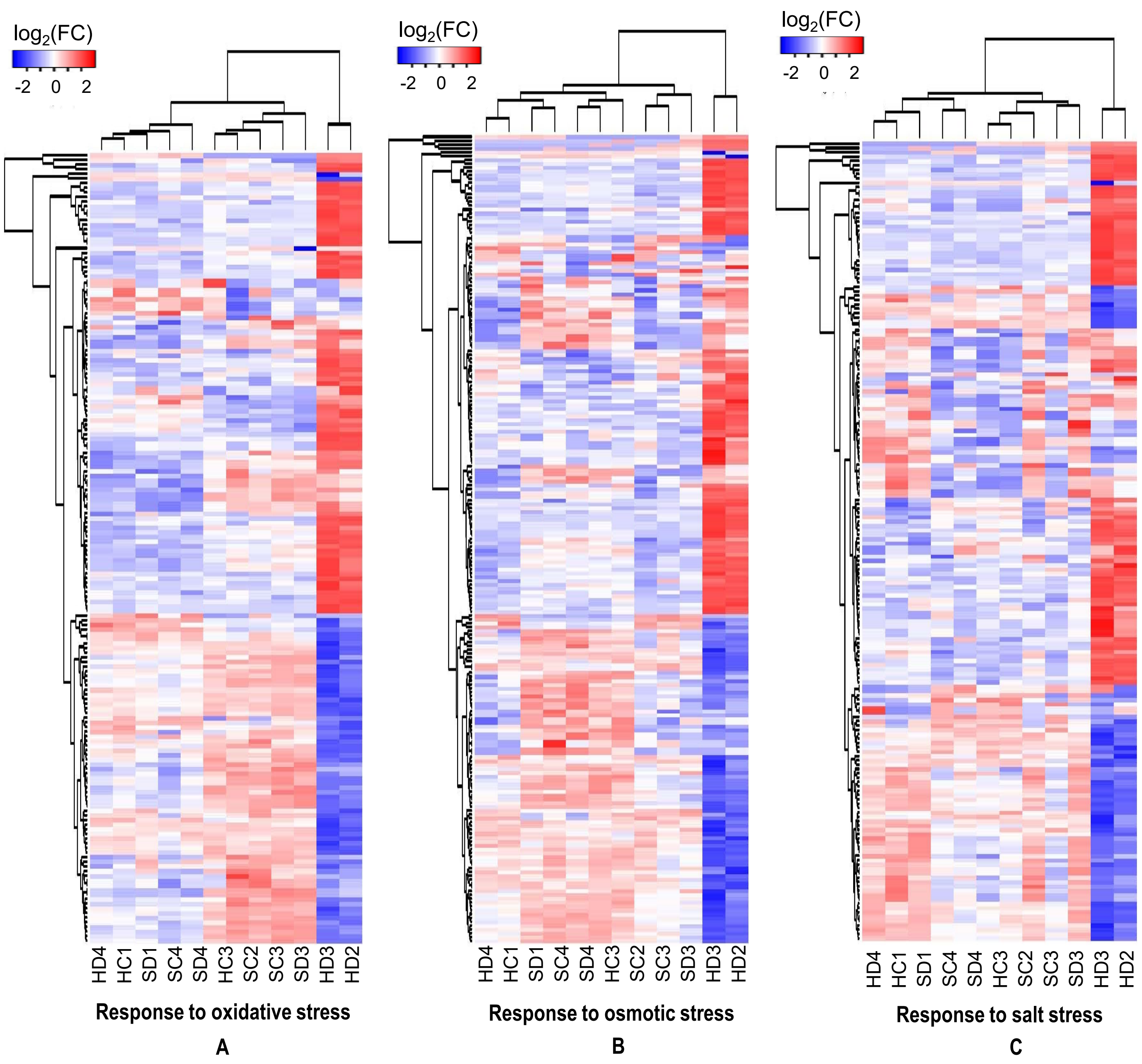


**Supplementary Figure 3**

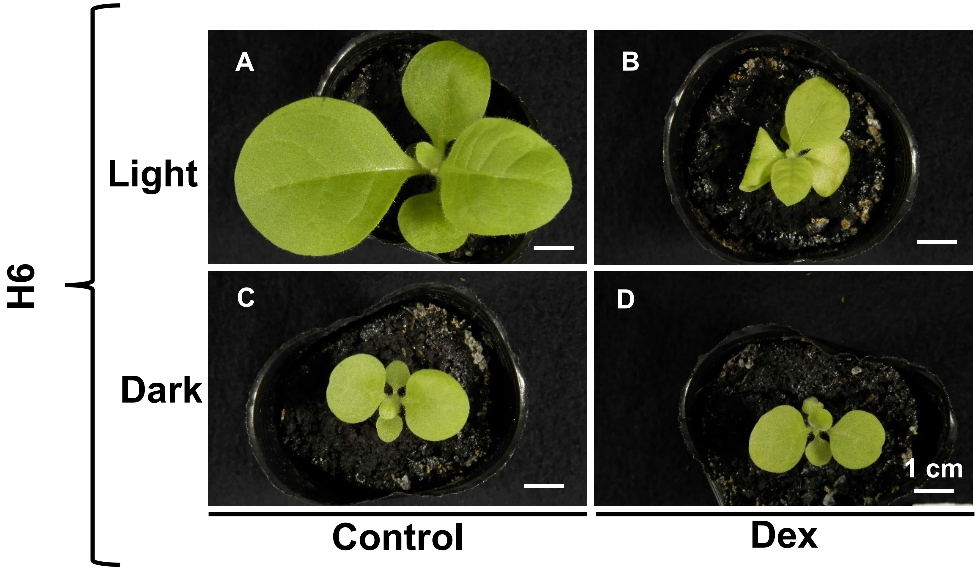


**Supplementary Figure 4**

**Table S2. Primers used for qRT-PCR analysis in the present study**

| Primer name | Sequence (5´→3´) |
|---|---|
| NtEF1α-qRT-867F | TGAGATGCACCACGAAGCTC |
| NtEF1α -qRT-917R | CCAACATTGTCACCAGGAAGTG |
| Nt-HSP90C-qRT-2111F | GGTTGAGCTCATCACCAT |
| Nt-HSP90C-qRT-2235R | CTTCTCCCTCTCATAAACTCC |
| Nt-ICS1-qRT-353F | CCACCCTCTCCAGCTCCTACT |
| Nt-ICS1-qRT-408R | TGGTCGGAACCAGGCAAT |
| NtLHCab-qRT-263F | ACCATCAAACTTGGAGAGATAC |
| NtLHCab-qRT-373R | GCCCATTCTTGAGCCTTTA |
| NtCHLI-qRT-94F | GCTTCTACACCCTTGTCTTC |
| NtCHLI-qRT-224R | ATTGGGACCTCCCTTTCT |

**Supplementary Table S3. Summary of RNA-Seq data and results of mapping the clean reads with *N. tabacum* TN90 reference transcriptome.**

| Samples | Raw Reads | Clean Reads | % Clean Reads | Mapped Reads | % of Mapped Reads | %GC | Clean Bases (GB) |
|---|---|---|---|---|---|---|---|
| HC 1 | 20750739 | 20578808 | 99.17 | 17026147.92 | 82.74 | 42 | 4.9 |
| HC 3 | 20740275 | 20608050 | 99.36 | 17694591.10 | 85.86 | 43 | 4.9 |
| HC 4 | 36041966 | 35624591 | 98.84 | 31677521.57 | 88.92 | 43 | 8.5 |
| HD 2 | 20455475 | 20329474 | 99.38 | 17600838.91 | 86.58 | 42 | 4.8 |
| HD 3 | 19697415 | 19536082 | 99.18 | 15658746.50 | 80.15 | 41 | 4.7 |
| HD 4 | 31430805 | 31240952 | 99.40 | 27430721.18 | 87.80 | 42 | 7.4 |
| SC 2 | 20017103 | 19680889 | 98.32 | 17027290.40 | 86.52 | 43 | 4.7 |
| SC 3 | 20290025 | 20171373 | 99.42 | 17639037.56 | 87.45 | 43 | 4.8 |
| SC 4 | 22178805 | 22058449 | 99.46 | 19887761.05 | 90.16 | 43 | 5.3 |
| SD 1 | 20550937 | 20409895 | 99.31 | 17754024.25 | 86.99 | 42 | 4.9 |
| SD 3 | 24776662 | 24512461 | 98.93 | 20121025.56 | 82.08 | 42 | 5.8 |
| SD 4 | 21715526 | 21547699 | 99.23 | 17654566.15 | 81.93 | 42 | 5.1 |
| | 278645733 (Total) | 276298723 (Total) | 99.17 (Average) | 237172272.10 (Total) | 85.60 (Average) | | |

**S, SR1 (non-transformant); H, line 4 of i-hpHSP90C; D, Dex treatment; C, Control.**
